# Genetic analysis of iris pigmentation in Swiss pig breeds identifies a missense *KITLG* variant as a potential causal factor for pale and heterochromatic irises

**DOI:** 10.1101/2025.09.02.673690

**Authors:** Wim Gorssen, Naveen Kumar Kadri, Negar Khayatzadeh, Alexander S. Leonard, Qiongyu He, Arnav Mehrotra, Stefan Neuenschwander, Hubert Pausch

**Affiliations:** Animal Genomics, Department of Environmental Systems Science, Universitätstrasse 2, 8092 Zürich, Switzerland; SUISAG, Allmend 10, Sempach, 6204, Switzerland; Animal Genetics unit, Tannenstrasse 1, 8092 Zürich

## Abstract

**Background:** Iris pigmentation is a heritable trait with a complex genetic architecture. While the genetic basis of iris pigmentation has been extensively studied in humans, little is known about iris pigmentation in pigs. Iris pigmentation in pigs varies from different shades of brown or pale irises to heterochromia manifesting either as different colors between both irises (heterochromia iridum) or multiple colors within a single iris (heterochromia iridis). This study investigates the genetics of iris pigmentation variability in the Swiss Landrace and Swiss Large White pig breeds.

**Results:** Iris pigmentation was phenotyped in 837 Swiss Landrace and 328 Swiss Large White pigs of which the majority also had array-derived genotypes. A high prevalence of heterochromia iridum (18.6%) was observed in the Swiss Landrace breed. Heritability estimates for iris pigmentation were high in both breeds (h² = 56.0–57.2%). Iris pigmentation was not genetically correlated with production traits. Genome-wide association analysis identified several loci associated with iris pigmentation, including regions near functional candidate genes such as *TYR, ALX4* and *DCT*. The strongest association was detected near the *KITLG* gene, which was identified as a candidate gene for iris pigmentation in a previous study on Italian Large White pigs. Fine-mapping identified a highly significantly associated (P=3.8×10^-11^) missense variant in *KITLG* (5_94084790_G>A, rs342599807, p.R124K) as a potential causal variant for pale and heterochromatic iris pigmentation in Swiss pigs.

**Conclusions:** Our findings provide new insights into the genetic architecture of iris pigmentation in pigs and indicate that *KITLG* plays a key role. The identification of a putative causal missense variant offers a foundation for further functional studies aiming to better understand pigmentation traits in pigs.

## Background

Eye color is determined by pigmentation of the iris, which is a two-layered compact structure composed of connective and smooth muscle tissue that regulates the amount of light entering the eye [1]). Iris pigmentation is determined by four main factors: (i) pigment granules in the posterior pigment epithelium, (ii) pigment concentration in stromal melanocytes, (iii) the type of melanin in these cells (brown-black eumelanin or red-yellow pheomelanin), and (iv) the light-scattering properties of the stromal matrix. Increased pigment in the iris stroma enhances light absorption, resulting in a darker iris pigmentation [1,2]. High melanin concentrations generally lead to (dark) brown irises, while low melanin concentrations lead to light-colored or pale irises whereas the absence of melanin (e.g. albinism) leads to red or pink irises. Heterochromia is the condition where an individual has two differently colored irises (heterochromia iridum or binocular heterochromia) or variations of color within a single iris (heterochromia iridis or sectorial heterochromia) [1].

Iris pigmentation is a heritable trait with a genetic architecture that is more complex than initially thought [3–7]. Heritability of iris pigmentation is very high in humans; in a twin study, Larsson et al. [8] estimated a broad sense heritability (H^2^) of 85% and a narrow-sense heritability (h^2^) of 51%. Simcoe et al. [7] found 124 independently associated QTL in 61 discrete genomic regions for iris pigmentation in a genome-wide association study (GWAS) involving almost 195,000 individuals.

The *tyrosinase*-encoding gene (*TYR*) has a large impact on iris pigmentation [9] as *tyrosinase* contributes to melanin synthesis, which can later be processed to both eumelanin and pheomelanin [6,10]. Other genes affecting iris pigmentation in humans are *oculocutaneous albinism II* (*OCA2*) and *HECT and RLD domain containing E3 ubiquitin protein ligase 2* (*HERC2)*, which are known to impact melanosome transport, and *melanocortin 1 receptor* (*MC1R*) which is a regulator of eumelanin and pheomelanin production [6]. Moreover, mutations in the *KIT ligand (KITLG)* gene, which also plays a role in melanogenesis [11], have been linked with skin, hair, and iris pigmentation in humans [12–15]. However, many other genes have been associated with iris pigmentation in humans (e.g., *TYRP*, *DCT*, *SLC24A4*, *SLC45A2*, *IRF4*) [7,16].

Only few studies have investigated iris pigmentation variability in pigs (Table 1). Moscatelli et al. [4] reported that 87% of Italian Large White pigs had brown pigmented irises (54.3% dark brown, 14.8% medium brown and 17.9% light brown), 3.8% had pale irises, 5.9% had heterochromia iridum and 3.2% had heterochromia iridis (2.3% unilateral versus 0.9 % bilateral). Nielsen and Lind [17] found 50% of Göttingen minipigs had brown irises, 47% had pale irises and 3% had heterochromia (type not specified). Gelatt et al. [3] found 78.5% of miniature pigs had brown irises, while 21.5% had some heterochromia pattern or pale irises. Moreover, they found an association between iris pigmentation and skin and hair color, where pigs with white skin and hair color had higher probability of generating offspring with pale irises or heterochromia, which was previously also observed by Yoshikawa [18]. Yoshikawa reported prevalences of 49.8% brown irises, 14.6% pale irises, 14.7% heterochromia iridum and 20.9% heterochromia iridis.

**Table 1.**
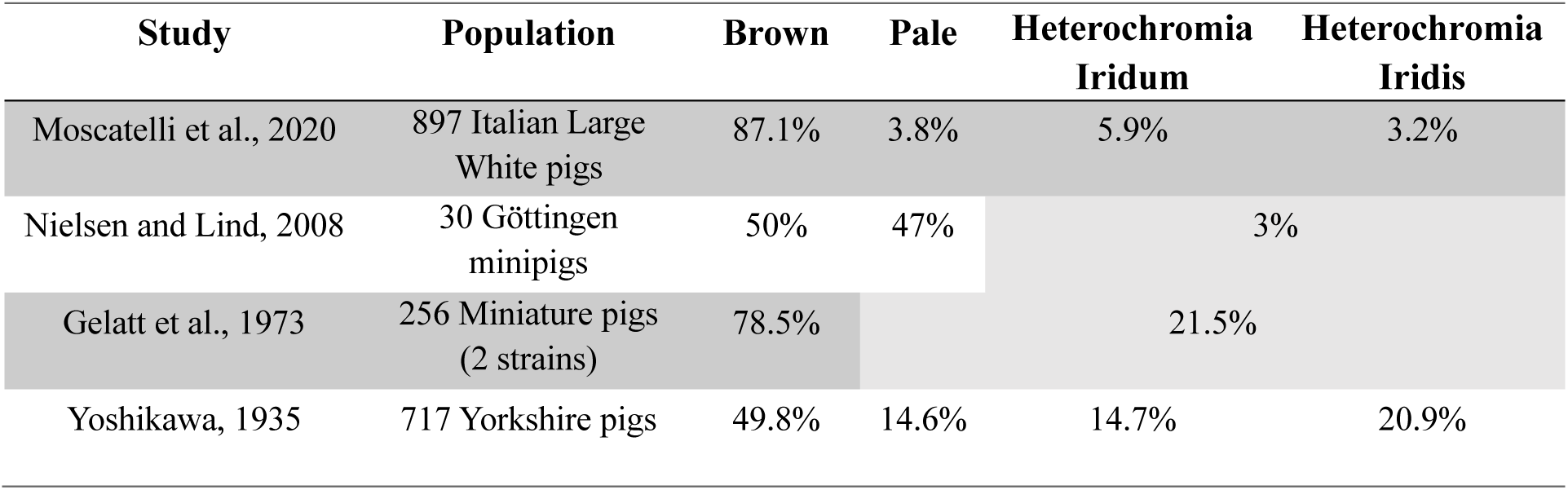
Prevalences of iris pigmentation in pigs reported by previous studies. Not all studies used a similar approach for phenotyping: Nielsen and Lind [17] did not differentiate between heterochromia iridum and iridis, while Gelatt et al. [3] did not differentiate between pale and heterochromia.

Moscatelli et al. [4] estimated that the h^2^ for heterochromia iridum versus brown irises was between 9.5% and 50.4% for different shades of brown irises in pigs. They also found a significant association (P<10^-6^) between several SNPs and iris pigmentation in a GWAS using medium-density microarray-derived genotypes. SNPs near the functional candidate gene *SLC45A2* located on the *sus scrofa* chromosome (SSC) 16 were associated with different shades of brown irises. Pale depigmented irises were associated with SNPs near *EDNRB* (SSC11) and *NOTCH2* (SSC4). Heterochromia iridis was associated with SNPs near *KITLG* (SSC5), *COL17A1* (SSC14) and regions on SSC6 and SSC8. Heterochromia iridum was associated with a region on SSC8.

Here, we investigate this variability in the Swiss Landrace and Swiss Large White pig breeds. First, the prevalence of five iris pigmentation traits is assessed in 1165 pigs. Second, heritabilities of iris pigmentation traits and their genetic correlations with production traits are estimated by integrating performance data, pedigree and genomic information in categorical animal models. Third, GWAS are carried out with haplotypes and imputed sequence variant genotypes to identify trait-associated variants. Last, potentially causal variants are identified in the Swiss Large White population using long and short read sequencing-derived variants.

## Methods

### Animal ethics statement

The pigs described in this study were kept in compliance with the Swiss legislations and were not raised or treated in any way for the purpose of this study.

### Animals and iris pigmentation collection

Iris pigmentation was assessed in 2024 in 1210 live pigs at the testing station of the breeding organization SUISAG and at five Swiss pig farms (SUISAG breeders). All pigs that were phenotyped at the SUISAG testing station were at the finishing stage, whereas pigs phenotyped at the farms were either gilts, sows or breeding boars. Iris pigmentation was assessed as described in Moscatelli et al. [4] (Table 2, Additional File 1 Figure S1). Each individual iris was scored as pale or one of three shades of brown: dark, medium or light brown. Differentiating between medium and dark brown irises was difficult under practical conditions with different light intensities, and so these two classes were later considered as ‘dark’. In the case of heterochromia iridis, the dominant iris color was noted first, followed by the secondary color. Some pigs, mainly Swiss Landrace individuals with drooping ears obscuring their eyes, could only be phenotyped for one iris. Animals with incomplete records (e.g., only one iris scored) were not considered for the downstream analyses. The final iris pigmentation dataset included 1165 pigs from the Swiss Landrace (N=837) and Swiss Large White (N=328) populations for which records for both irises were available. The Swiss Large White animals descend from either a dam or a sire line, which trace back to the same ancestral population but were bred independently since 2002 [19]. Due to a relatively recent common ancestral population, we consider the Swiss Large White sire and dam line as one population but account for population stratification and relationships in all statistical models (below).

**Table 2.** Overview of iris pigmentation phenotypes used in this study.

| **Iris pigmentation phenotype** | **Description** |
| --- | --- |
| **Dark brown** | Pigs with both irises completely medium or dark brown |
| **Light brown** | Pigs with both irises completely light brown |
| **Pale** | Pigs with both irises completely pale/depigmented |
| **Heterochromia iridum** | Pigs with one iris completely pale/depigmented and the other iris completely brown of any shade |
| **Heterochromia iridis** | Pigs with one or both irises with both pale/depigmented and any shade of brown within the same iris<br>Unilateralis: one iris affected<br>Bilateralis: both irises affected |

### Phenotype data for production traits

Phenotypes for daily gain (g/d), backfat depth (mm) and loin muscle depth (mm) were available for 611 Swiss Landrace and 283 Swiss Large White pigs. These phenotypes were either recorded on live animals at the testing station of SUISAG or during a field test at the breeder farm. These data were visually inspected for outliers and normality via histograms (Additional File 2 Figure S2, Additional File 3 Figure S3), but no outliers were detected or removed.

### Genotype data

Microarray-derived genotypes were provided by SUISAG. The genotype data comprised 77,056 SNPs and 1164 pigs before quality control (757 Swiss Landrace, 407 Swiss Large White). The phenotypic and genotypic datasets overlapped largely, but some animals had only phenotype or only genotype records. The coordinates of the SNPs were determined according to the current porcine reference genome (*Sscrofa11.1*). Reference and alternate alleles were updated accordingly for each SNP and SNPs with duplicated positions were removed. Only autosomal SNPs were retained (71,915 SNPs). We used PLINK v1.9 [20] to exclude 32 Swiss Landrace and 13 Swiss Large White individuals with call-rate below 95% or an excess of heterozygosity (i.e., deviating more than 3 standard deviations from the mean). We excluded SNPs with call rate lower than 95%, minor allele frequency below 1% and SNPs deviating significantly from Hardy-Weinberg proportions (P<0.0001). The SNP-based quality control was performed for each breed separately. The final genotype dataset contained 725 pigs and 56,096 SNPs for Swiss Landrace and 394 pigs and 55,776 SNPs for Swiss Large White. A combined dataset was created based on the overlapping SNPs after per breed QC, containing 1119 pigs and 49,755 SNPs. Of the genotyped individuals, phenotypic records were available for 586 Swiss Landrace pigs, 303 Swiss Large White pigs and 889 pigs in the combined dataset.

After quality control, a principal component analysis (PCA) was performed with PLINK v1.09 (-pca) on the combined dataset to examine population structure (Additional File 4 Figure S4).

### Phasing and imputation

The array-derived genotypes were phased, and sporadically missing genotypes were imputed for each breed separately using Beagle v5.4 software [21] (beagle.01Mar24.d36.jar). The phased array-derived genotypes of Swiss Large White pigs were imputed to the sequence level based on an in-house reference panel [22] consisting of 120 Swiss Large White pigs that were sequenced (Illumina short-read) to at least 10x autosomal coverage (mean=15.9; sd=5.0; max=36.8) and which had genotypes at 31,652,340 autosomal sequence variants. None of the sequenced pigs had iris pigmentation phenotypes. The genetic relationship between the genotyped population and the sequenced reference population was inspected through PCA (Additional File 5 Figure S5). Reference-based imputation was performed using Beagle with an effective population size of 100, a window of 10, overlap 2, and burnin 5 [23].

Sequence variant genotypes for the Swiss Landrace pigs were inferred using the SWine IMputation haplotype reference panel (SWIM; https://www.swimgeno.org/) [24].

After imputation, there were 31,652,340 SNPs with a mean r^2^ of 0.710 for Swiss Large White and 34,615,361 SNPs with a mean r^2^ of 0.725 for Swiss Landrace. However, only SNPs with a minor allele frequency > 0.01 and an r^2^ value > 0.5 were retained, leading to 22,378,549 SNPs with mean r^2^ of 0.941 for Swiss Large White and 15,631,491 SNPs with mean r^2^ of 0.935 for Swiss Landrace. For the combined analysis of the Swiss Large White and Swiss Landrace pigs, overlapping SNPs after per-breed QC were retained (12,116,974 SNPs).

### Estimation of genetic parameters

Genetic parameters were estimated with *gibbsf90+* software [25] using a Bayesian method implementing single-step genomic prediction (OPTION SNP_file). A threshold model was used with five categorical phenotypes (OPTION cat 5; dark brown, light brown, pale, heterochromia iridum and heterochromia iridis) for each breed separately. All models were run with 500,000 samples, a burnin of 100,000 and a sample rate of 1,000. Mean estimates and 95% highest posterior density intervals (HPD) were reported.

The estimated sire-dam models were of the form:

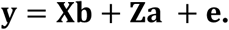

Here, **y** is the vector with categorical iris pigmentation phenotypes; **b** is a vector with fixed effects and covariates. The fixed effect was farm-date (6 levels); **a** is a vector containing additive genetic animal effects (2496 animals in the pedigree and 725 with genotype information for Swiss Landrace; 3710 animals in the pedigree and 394 with genotype information for Swiss Large White). The assumption is that **a** follows a normal distribution for the **H** matrix following [26–28], using single-step genomic evaluation with both pedigree (**A**) and genomic (**G**) relationship matrices: 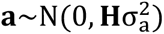. **e** is the vector of residual effects assumed to follow a normal distribution 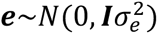; **X** and **Z** are incidence matrices for respectively fixed effects and random animal effects. The **H** matrix was constructed using the array-derived medium density genotypes.

Next, we estimated genetic correlations (r_g_) of iris pigmentation with the production traits life daily gain (g/day), backfat thickness (mm) and loin muscle depth (mm) using bivariate models. Genetic correlations were only retained for the Swiss Landrace population, as sample size was insufficient to reliably estimate genetic correlations for our Swiss Large White dataset (N=328). Genetic correlations were estimated as follows:

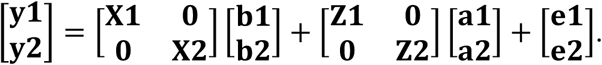

Similar to the five-trait sire-dam model, **y1** represents a vector with categorical iris pigmentation phenotypes with 5 categories and **y2** represents a vector with one of the continuous production trait phenotypes; **b1** and **b2** are vectors containing fixed effects and covariates, which were farm-date of sampling (6 levels), sex (3 levels: sow, boar, barrow) and age at recording of production data (covariate); **a1** and **a2** are vectors of additive animal genetic effects, assumed to follow a normal distribution for the **H** matrix using single-step genomic evaluation:

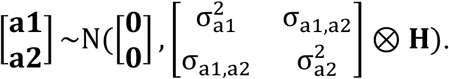

**e1** and **e2** are vectors of residual effects, assumed to follow a normal distribution 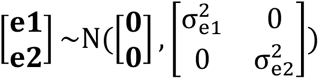; **X1**, **X2**, **Z1** and **Z2** are incidence matrices for fixed effects and random animal effects.

### Genome wide association study

Genome wide association studies were conducted separately for the combined dataset and for both breeds at the haplotype level and on the imputed sequence level. Twelve case-control scenarios of contrasting iris pigmentation traits were considered (Table 3). In total, 72 GWAS analyses were performed: twelve scenarios for three populations at the haplotype and the imputed sequence level.

**Table 3.**
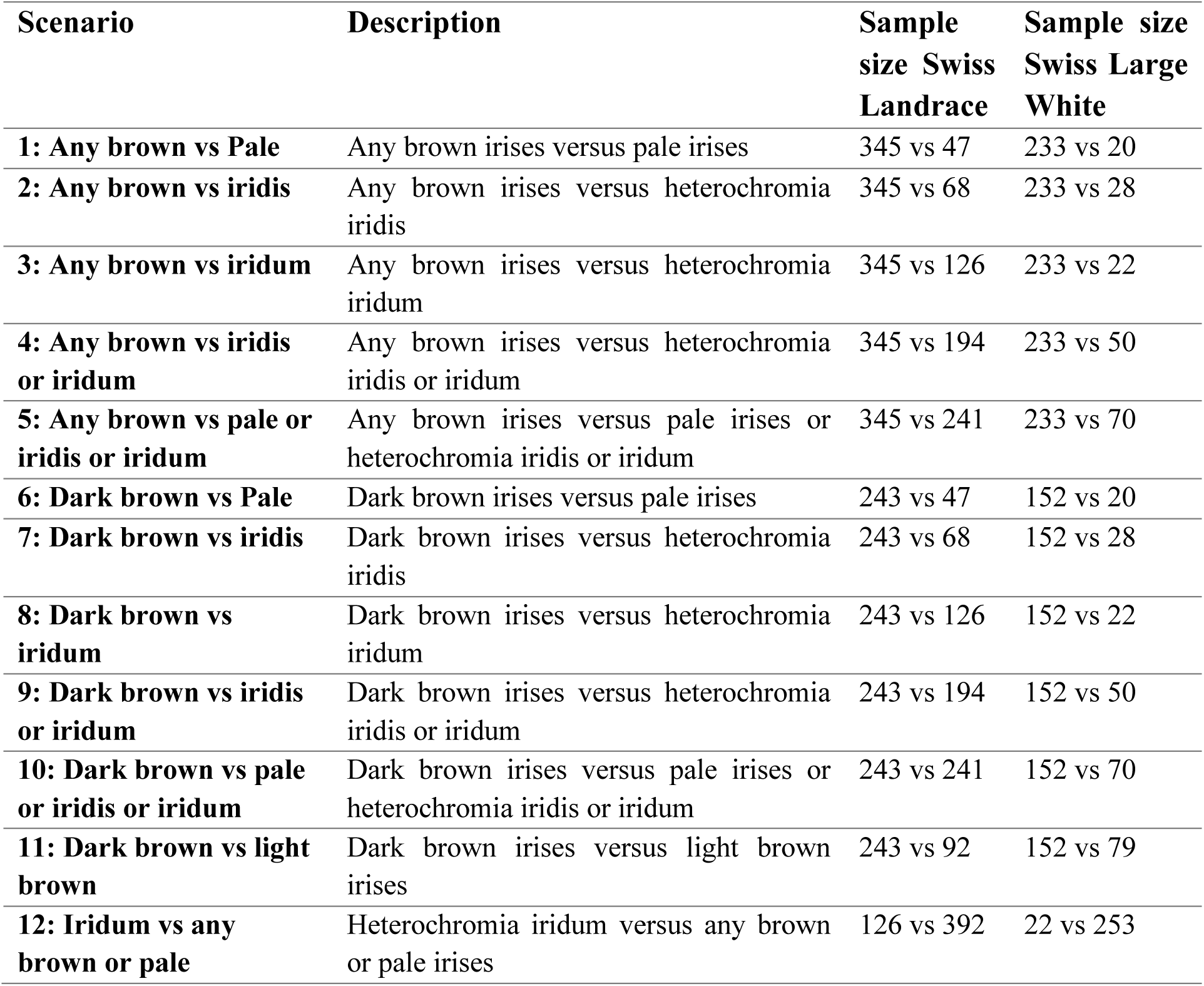
Case-control contrast scenarios for iris pigmentation used for GWAS analyses. The sample size for the combined analysis of Swiss Landrace and Swiss Large White is the sum of the sample sizes of both breeds.

We used the GCTA software (gcta-1.94.1) [29] to conduct GWAS with imputed sequence variants considering the genomic relationship matrix and its first four principal components to account for population stratification (--mlma, --grm, --qcovar).

Haplotype-based association analysis was conducted using a sliding window of 10 SNPs with an overlap of 5 SNPs, while considering the first twenty principal components and a minor allele frequency cutoff of 1%. For haplotypes segregating at a frequency between 0.01 and 0.99, difference in its count in case and control groups was tested using Fisher’s exact test.

Manhattan plots were generated with the *manhattan* function of the *qqman* package in R [30] using a genome-wide significance threshold of P<5×10^-8^ and a suggestive significance threshold of P<1×10^-5^.

### Identification of candidate genes

Genes annotated in the Sscrofa11.1 genome assembly [31] within a ±2 Mb region surrounding all significantly associated SNPs and haplotypes were identified by downloading annotated genes using the biomaRt package in R for Ensembl database version 114 [32]. A zoomplot was made via an in-house R-script for genomic regions associated with iris pigmentation, implementing information on linkage disequilibrium and imputation accuracy (example in Figure 2D). Positional candidate genes were subsequently evaluated for their potential influence on the associated phenotypes based on a comprehensive review of relevant literature.

### Functional annotation of associated variants

Ensembl’s Variant Effect Predictor (VEP, version 114.0) tool [33] was used to predict functional consequences for all variants with P<10^-5^ in any of the GWAS scenarios (Table 3). The *Sus scrofa* reference genome (*Sscrofa11.1*) and the corresponding Ensembl transcript annotations were considered. Predicted variant effects were categorized according to sequence ontology terms (e.g., synonymous_variant, missense_variant, intron_variant). Variants were also annotated with gene symbols, transcript information, regulatory region consequences, and functional impact scores (SIFT and PolyPhen) where available.

### Structural variants near *KITLG* gene

The presence of structural variants was investigated in the window from 92 to 96 Mb on chromosome 5. Within our sequenced cohort of 120 Swiss Large White pigs, we assessed the haplotype status for the most significantly associated haplotype in this region. We defined three groups: homozygous haplotype carrier, heterozygous haplotype carrier and no haplotype carrier. Using Mosdepth [34], the average normalized (z-score) coverage per 250 bp window was calculated and compared between groups. Windows with an absolute difference >3 (3 standard deviations from mean) were inspected via Integrative Genomics Viewer (IGV_2.19.5) [35].

To identify structural variants associated with the haplotype, we also analyzed 18 HiFi-sequenced samples, including 10 Swiss Large White pigs and 8 hybrid animals with a Swiss Large White parent. High molecular weight DNA was extracted from frozen ear or tail tissue or blood, all provided by SUISAG for unrelated studies, using the NEB Monarch® HMW DNA Extraction Kit. Sequencing was performed on the PacBio Sequel IIe platform, with one SMRT cell per sample. HiFi reads were aligned to the same reference genome using pbmm2 v1.17 [36] followed by structural variant calling with sawfish v1.0.1 [37].

## Results

### Iris pigmentation distribution and prevalence per breed

Two thirds (65%) of 837 Swiss Landrace pigs and three quarters (76.5%) of 328 Swiss Large White pigs had brown irises, whereas 7.3% and 6.4% had pale irises (Table 4, Figure 1). The prevalence of heterochromia iridum differed considerably between breeds (18.6% in Swiss Landrace pigs compared to 7.0% in Swiss Large White pigs). A high prevalence of heterochromia iridum was observed in all six Swiss Landrace farms (minimum 13.6% to maximum 23.4%; Additional File 6 Table S1). Heterochromia iridis had a prevalence of 9.0% in Swiss Landrace versus 10.0% in Swiss Large White.

**Figure 1.**
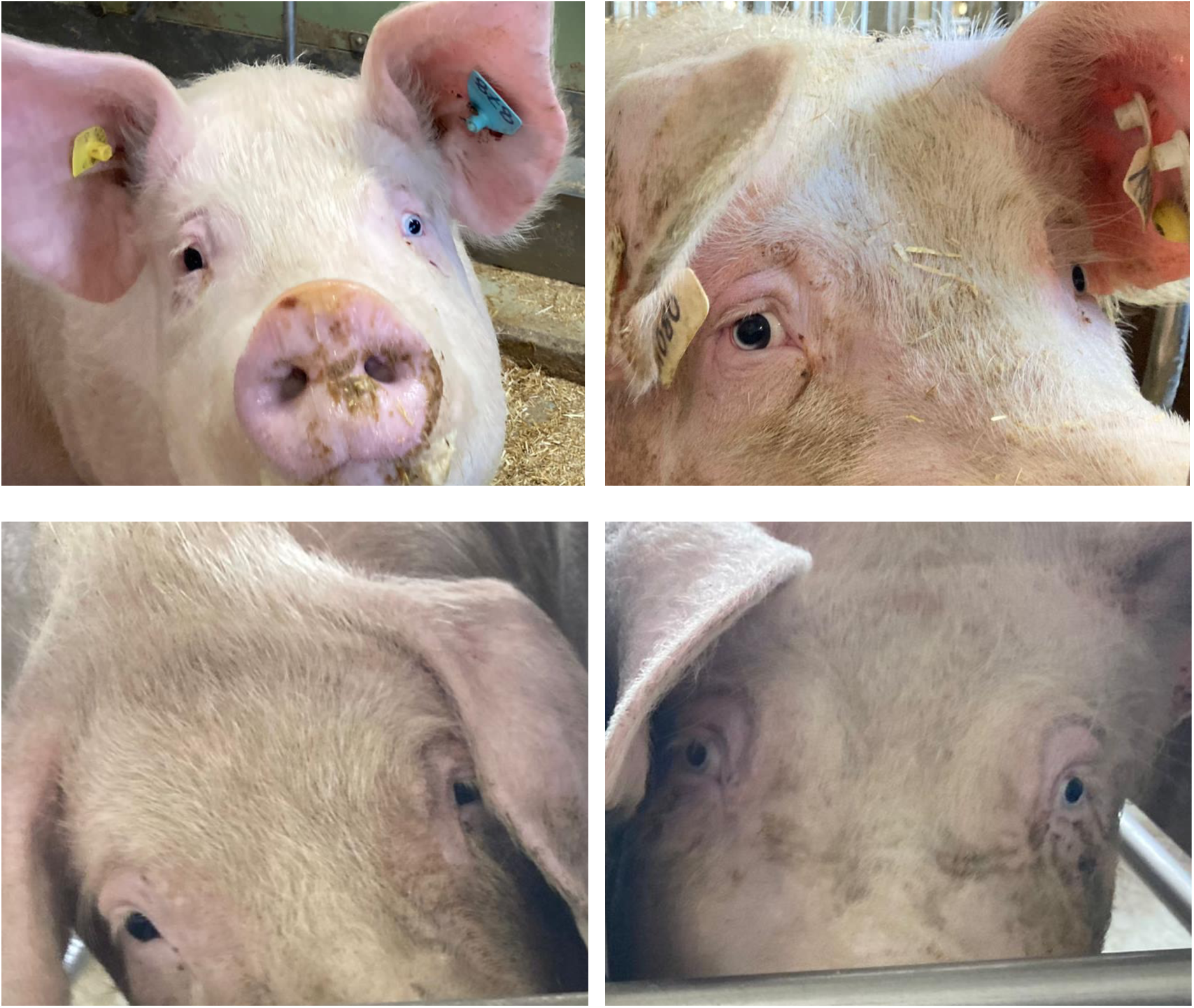
Variability in iris pigmentation observed in Swiss Landrace pigs. Top left: heterochromia iridis, with a brown colored right iris and a pale colored left iris. Top right: Heterochromia iridis unilateralis, where the right iris is both brown and pale. Bottom: two sows with different iris pigmentation: left, a sow with brown pigmented irises, right, a sow with pale pigmented irises.

**Table 4.** Summary of observed iris pigmentation phenotypes in Swiss pig breeds. In total, there were 837 Swiss Landrace pigs and 328 Swiss Large White pigs with an iris pigmentation phenotype. The number of animals with a given iris pigmentation phenotype is shown and the relative percentage within the breed is given between brackets. A detailed overview per farm and per breed is given in Additional File 6 Table S1.

| **Iris pigmentation phenotype** | **Swiss Landrace**<br>**(N=837)** | **Swiss Large White**<br>**(N=328)** |
| --- | --- | --- |
| **Dark Brown** | 407 (48.6%) | 166 (50.6%) |
| **Light Brown** | 137 (16.4%) | 85 (25.9%) |
| **Pale** | 61 (7.3%) | 21 (6.4%) |
| **Heterochromia Iridum** | 156 (18.6%) | 23 (7.0%) |
| **Heterochromia Iridis Unilateralis** | 64 (7.6%) | 28 (8.5%) |
| **Heterochromia Iridis Bilateralis** | 12 (1.4%) | 5 (1.5%) |

Sows with dark brown irises had the highest incidence of offspring with dark brown irises (64.1%), whereas sows with pale irises had a low incidence of offspring with dark brown irises (38.2%) (Additional File 7 Table S2). Moreover, sows with heterochromia iridum had the highest incidence of offspring with heterochromia iridum (25.0%).

### Genetic parameters

The h^2^ of iris pigmentation was 57.2% (HPD: 24.2 to 78.4%) in Swiss Landrace and 56.0% (HPD: 15.6 to 90.9%) in Swiss Large White (Table 5) on the liability scale. The genetic correlations between iris pigmentation and the three production traits studied had HPD intervals that included zero, indicating no strong evidence for a non-zero correlation.

**Table 5.**
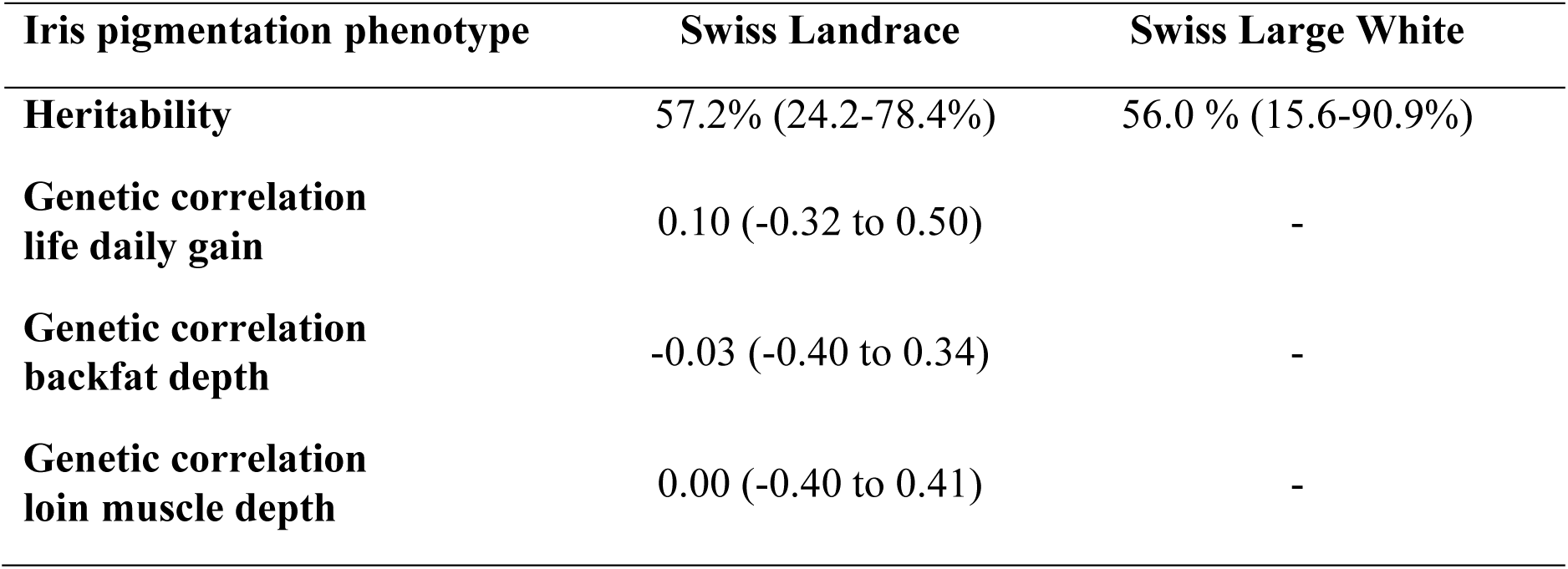
*Heritability and* genetic correlation estimates for iris pigmentation and production traits. Heritability estimates are on the liability scale. The 95% highest posterior density estimates (HPD) are given between brackets. It should be noted that the HPD intervals are large, due to the limited sample size. Genetic correlations were not estimated for Swiss Large White due to insufficient sample size (N=328). Detailed estimates of variance components are given in Additional File 8 Table S3 and Additional File 9 Table S4.

### Genome wide association study for iris pigmentation

Association tests using haplotypes and imputed sequence variants were conducted to identify genomic regions associated with iris pigmentation. In total, 72 association studies were performed using both SNP-based and haplotype-based approaches. The genomic inflation factor (λ) averaged 1.18 (SD = 0.34; range: 0.88–2.20), suggesting that population stratification and related confounding were generally well controlled although inflation was apparent for some association studies. Several regions were significantly associated with iris pigmentation at the genome-wide threshold (P<5×10^-8^) across the twelve GWAS scenarios (Table 6, Figure 2).

**Table 6.** Overview of genetic regions significantly associated (P<5×10^-8^) across different GWAS scenarios. The results are ordered by chromosome number and location. Per 1 Mb bin, the most significantly associated SNP is shown per GWAS scenario and per breed. For haplotype-based analyses, no top associated variant was given, as this is uninformative. SSC: Sus scrofa chromosome. Freq: Minor allele frequency or haplotype frequency. Detailed estimates including odds ratios and effect sizes for all significant variants are provided in Additional File 11 Table S6.

| Case-control GWAS scenario | Breed | SSC | Top variant or haplotype (bp) | Method | Freq | P-value | Candidate gene |
| --- | --- | --- | --- | --- | --- | --- | --- |
| Dark brown vs pale or iridis or iridum | Landrace | 2 | 12623444-12799159 | Haplotype | 0.42 | $4.5 \times 10^{-8}$ | |
| Any brown vs iridis or iridum | Landrace | 2 | 13406605-14417730 | Haplotype | 0.39 | $3.4 \times 10^{-8}$ | |
| Any brown vs pale or iridis or iridum | Landrace | 2 | 13406605-14417730 | Haplotype | 0.39 | $1.8 \times 10^{-8}$ | |
| Any brown vs pale or iridis or iridum | Landrace | 5 | 94040632-94284630 | Haplotype | 0.16 | $8.5 \times 10^{-9}$ | *KITLG* |
| Any brown vs pale or iridis or iridum | Large White | 5 | 94037661-94216956 | Haplotype | 0.39 | $1.8 \times 10^{-8}$ | *KITLG* |
| Any brown vs iridis | Landrace | 5 | 94178087 | Imputed | 0.23 | $4.8 \times 10^{-9}$ | *KITLG* |
| Dark brown vs iridis | Landrace | 5 | 94178087 | Imputed | 0.24 | $4.2 \times 10^{-8}$ | *KITLG* |
| Any brown vs pale or iridis or iridum | Large White | 5 | 94089673 | Imputed | 0.43 | $1.3 \times 10^{-9}$ | *KITLG* |
| Dark brown vs iridis or iridum or pale | Large White | 5 | 94089673 | Imputed | 0.45 | $1.3 \times 10^{-8}$ | *KITLG* |
| Any brown vs pale or iridis or iridum | Combined | 5 | 94242738 | Imputed | 0.26 | $2.1 \times 10^{-12}$ | *KITLG* |
| Any brown vs iridis or iridum | Combined | 5 | 94242738 | Imputed | 0.25 | $2.3 \times 10^{-10}$ | *KITLG* |
| Any brown vs iridis | Combined | 5 | 94242738 | Imputed | 0.24 | $4.4 \times 10^{-10}$ | *KITLG* |
| Dark brown vs pale or iridis or iridum | Combined | 5 | 94259064 | Imputed | 0.26 | $3.0 \times 10^{-10}$ | *KITLG* |
| Dark brown vs iridis | Combined | 5 | 94495591 | Imputed | 0.23 | $4.7 \times 10^{-9}$ | *KITLG* |
| Dark brown vs iridis or iridum | Combined | 5 | 94259064 | Imputed | 0.25 | $4.7 \times 10^{-9}$ | *KITLG* |
| Any brown vs pale | Large White | 8 | 10799489 | Imputed | 0.06 | $3.1 \times 10^{-9}$ | |
| Any brown vs pale | Large White | 10 | 7921953 | Imputed | 0.06 | $2.2 \times 10^{-8}$ | |
| Dark brown vs pale | Large White | 10 | 28724329 | Imputed | 0.02 | $2.2 \times 10^{-8}$ | |
| Iridum vs any brown or pale | Large White | 11 | 62703128 | Imputed | 0.02 | $7.2 \times 10^{-10}$ | *DCT* |
| Any brown vs iridum | Large White | 11 | 62703128 | Imputed | 0.02 | $4.2 \times 10^{-9}$ | *DCT* |
| Dark brown vs iridum | Large White | 11 | 62703128 | Imputed | 0.02 | $2.2 \times 10^{-8}$ | *DCT* |
| Any brown vs iridum | Large White | 11 | 63954396 | Imputed | 0.01 | $2.7 \times 10^{-8}$ | *DCT* |
| Iridum vs any brown or pale | Large White | 11 | 63954396 | Imputed | 0.01 | $3.9 \times 10^{-9}$ | *DCT* |
| Iridum vs any brown or pale | Large White | 11 | 65138862 | Imputed | 0.01 | $3.9 \times 10^{-9}$ | *DCT* |
| Any brown vs iridum | Large White | 11 | 65138862 | Imputed | 0.01 | $2.7 \times 10^{-8}$ | *DCT* |
| Any brown vs pale | Large White | 16 | 73678712 | Imputed | 0.02 | $8.8 \times 10^{-10}$ | |
| Any brown vs iridis | Landrace | 17 | 25089995 | Imputed | 0.01 | $2.5 \times 10^{-8}$ | |

**Figure 2.**
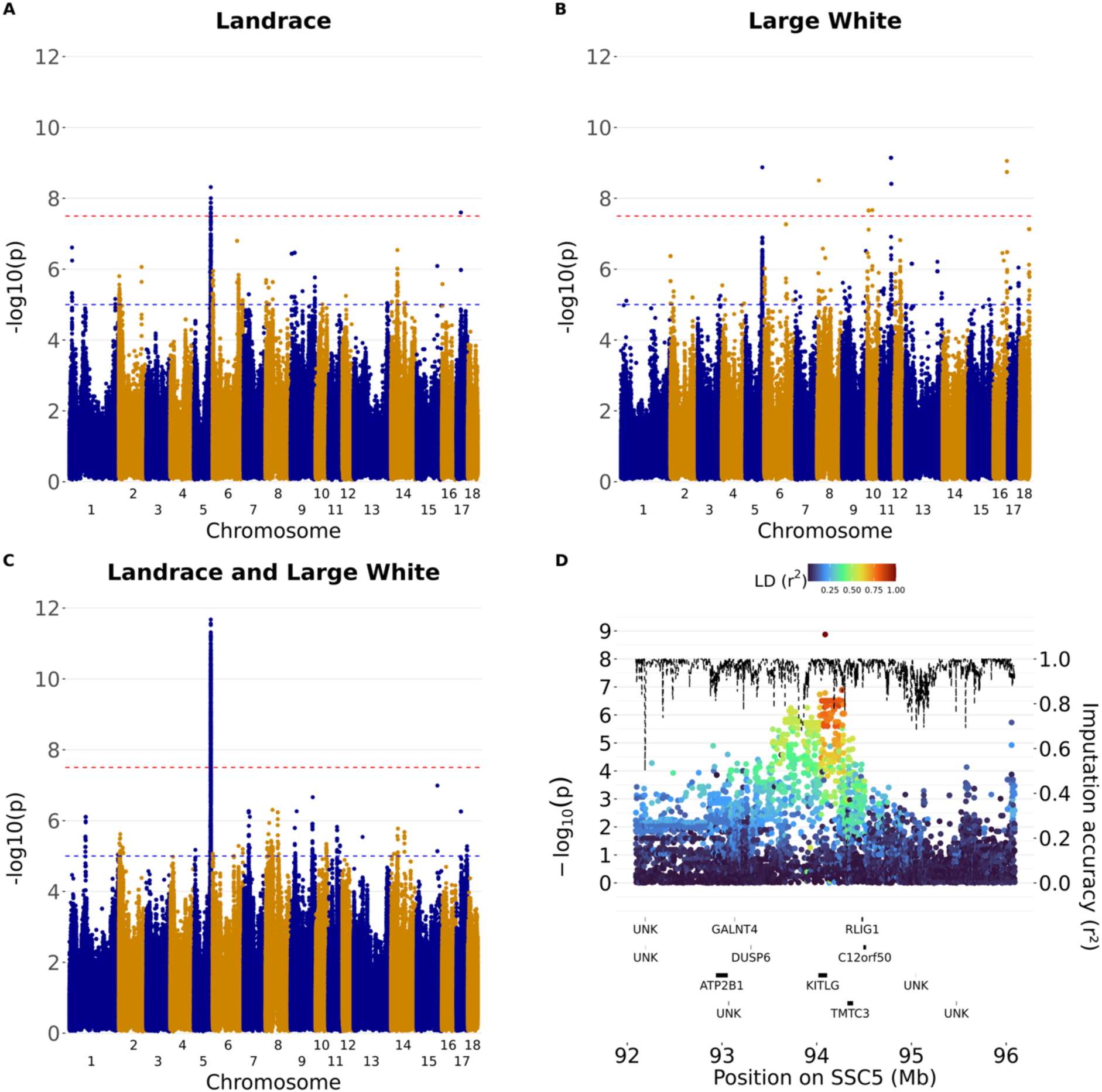
Manhattan plots for iris pigmentation in Swiss pig breeds. For each variant, the most significant association was kept over all twelve GWAS scenarios (Table 3) for Swiss Landrace (A), Swiss Large White (B), and the combined analysis (C). Panel D shows the zoomplot for the association of any brown irises versus pale or heterochromia iridis or iridum (scenario 5, Table 3) in Swiss Large White. The negative logarithm of the p-value is given on the left y-axis, the chromosomal location in Mb is given on the x-axis. For the Manhattan plots, the red horizontal line indicates the genome-wide threshold of P<5×10-8, while the blue horizontal line indicates the suggestive threshold of P<1×10-5. For the zoomplot, imputation accuracy is given as a dashed line (right y-axis) and the linkage disequilibrium (r^2^) with the top variant is colored according to the legend. Results per GWAS scenario are provided in Additional File S10 Table S5 and Additional File S11 Table S6.

Across all combinations tested, the strongest associations between iris pigmentation and genetic markers were found on SSC5 between 94,037,661 bp and 94,495,591 bp encompassing the *KITLG* gene (Table 6, Figure 2). This region was identified in both breeds, and in the multi-breed analysis. Significant markers or haplotypes were detected in the case-control scenarios 2, 4, 5, 7, 8, 9, and 10 (see Table 3), in which animals with brown or dark brown eyes are cases, and animals with pale, iridis and/or iridum eyes are controls. Across all GWAS conducted, the most significant association was detected in the imputed SNP-based analysis of scenario 5 (Chr5:94,242,738; P=2.1×10^-12^).

Significant associations other than those within the interval from 92 Mb to 96 Mb on SSC5 were only found within but not across breeds. Significant trait-variant associations were observed in the Swiss Landrace population on SSC2 and SSC17. On SSC2, the most significantly associated haplotypes were found for scenario 10 (Chr2:12,623,444-12,799,159; P= 4.5×10^-8^) and scenarios 4 and 5 (Chr2:13,406,605-14,417,730; P=3.4×10^-8^ and P=1.8×10^-8^ respectively). On SSC17, the most significant association was detected in the sequence-based analysis of scenario 2 (Chr17:25,089,995; P=2.5×10^-8^).

In the Swiss Large White population, we identified significant associations on SSC8, SSC10, SSC11 and SSC16. On SSC8, the most significant association was detected in the sequence-based analysis of scenario 1 (Chr8:10,799,489; P=3.1×10^-9^). On SSC10, significant variants were detected in scenario 1 (Chr10:7,921,953; P=2.2×10^-8^) and scenario 6 (Chr10:28,724,329; P=2.2×10^-8^). On SSC11, multiple significant variants were detected between 62,703,128 bp and 65,138,862 bp, encompassing the functional candidate gene *DCT*. Significant SNPs or haplotypes were detected in the case-control scenarios 3, 8, and 12 with the most significant association for scenario 12 (Chr11:62,703,128; P=7.2×10^-10^). On SSC16, there was a significantly associated variant for scenario 1 (Chr16:73,678,712; P=8.8×10^-10^).

An overview of the 536 haplotypes and 14,835 SNPs surpassing the suggestive threshold (P<1×10^-5^) is given in both Additional File 10 Table S5 and Additional File 11 Table S6. Although these associations are not all significant on the genome-wide threshold and may contain false-positives, manual inspection of annotated genes surrounding the top variants was conducted. Upon comparing associations across the Swiss Landrace and Swiss Large White populations, overlapping regions with associations (P<1×10^-5^) are independently present in both breeds on SSC2, SSC5 (discussed above), SSC6, SSC8 and SSC15.

On SSC2, overlapping associations between breeds were found in two regions: in the region between 7.6 Mb and 7.8 Mb for scenarios 1 and 2 and in the region between 17.8 Mb and 18.9 Mb encompassing the functional candidate gene *ALX4* for scenarios 1, 3, 5, 6, 10 and 12. Within these regions, the most significant association was found for the haplotype analysis for scenario 5 (Chr2:17,821,105-18,078,630; P=3.4×10^-7^). On SSC6, both breeds had associations (P<10^-5^) between 2.8 Mb and 3.6 Mb for scenarios 2, 3, 8 and 12, with the strongest association for a variant in scenario 2 (Chr6:2,872,204; P=9.6×10^-7^). On SSC8, associations were detected in the region of 40.7 Mb to 42.9 Mb encompassing the *KIT* gene for scenarios 5, 9 and 10. The strongest association was found for an imputed sequence variant within the Swiss Landrace population for scenario 10 (Chr8:40,778,203; P=2.3×10^-6^). On SSC15, overlapping associations were found in the region between 114.8 Mb and 115.3 Mb for scenarios 1, 5 and 10. The strongest association was found for a haplotype in scenario 5 (Chr15:114,816,163-115,237,432; P=2.3×10^-6^).

A sequence variant (rs333452789, Chr9:22,593,530) was suggestively associated (P=3.4×10^-7^) in scenario 1 in the Swiss Landrace population. This variant is in an intron of the functional candidate gene *TYR*.

### Functional annotation of variants associated with iris pigmentation

To functionally annotate variants associated with iris pigmentation, we used Ensembl’s VEP on 14,835 variants that were associated (P<10⁻⁵) in at least one case-control scenario (Additional File 12 Table S7). Among them, 90.1% were known variants, and 9.9% were novel. A total of 305 genes and 214 regulatory features were overlapped by these variants. Most annotated variants were in intronic (51%) and intergenic (32%) regions, followed by downstream (6%) and upstream (5%) gene regions.

Only a small proportion affected coding regions directly, with 27 synonymous variants and 25 missense variants. One high-impact frameshift variant was detected (2_825376_A>AC), although not in a known gene and only with an association of P=9.3×10^-6^ in GWAS scenario 1 for Swiss Large White pigs. Moderate-impact coding variants (missense) were identified in several genes, including *KITLG*, *CEP290*, *TYR*, *APCDD1L*, *ANKRD60*, and *RAB22A*. From the variants in coding regions, only a missense variant in *KITLG* (5_94084790_G>A; rs342599807, KITLG:p.R124K; most significant association of P=3.8×10^-11^ for scenario 5 in the combined analysis and imputation r^2^=1) and a missense variant in *CEP290* (5_94453633_C>T; rs334726982, CEP290:p.A1958V; most significant association of P=3.4×10^-9^ for scenario 3 in the combined analysis and imputation r^2^=1), surpassed the genome-wide significance threshold.

### Structural variants near *KITLG* gene

We examined the interval from 92 Mb to 96 Mb on SSC5 encompassing *KITLG* for the presence of structural variants in Swiss Large White pigs, as several strongly associated haplotypes and variants within this interval indicate that this region has an impact on iris pigmentation in that population. We focused on the haplotype which was most strongly associated (P= 1.8×10^-8^; Table 6) with iris pigmentation in the genotyped Swiss Large White pigs. This haplotype consists of 10 consecutive SNPs and spans from 94,037,661 bp to 94,216,956 bp, thereby partly overlapping the *KITLG* gene (Additional File 10 Table S5, Additional File 11 Table S6). This haplotype occurs at a frequency of 21.2% in the sequenced cohort of 120 Swiss Large White individuals and is also in perfect linkage disequilibrium (r=1.00) with the *KITLG* missense variant 5_94084790_G>A. Our association tests indicate that haplotype carriers have an increased prevalence of pale or heterochromatic irises. Within the sequenced cohort, there were 6 homozygous haplotype carriers, 39 heterozygous carriers, and 75 non-carriers. We identified 8 regions for which the normalized sequence coverage differed by a factor of at least 3 between homozygous haplotype carriers and non-carriers possibly indicating the presence of deletions or duplications (Chr5:93,895,250-93,895,500; Chr5:93,956,750-93,959,250; Chr5:94,015,000-94,015,250; Chr5:94,036,000-94,036,250; Chr5:94,046,750-94,047,250; Chr5:94,282,250-94,282,500; Chr5:94,286,000-94,286,250; Chr5:94,307,750-94,308,000). For all these regions, the normalized sequence coverage was low for homozygous haplotype carriers, intermediate for heterozygous carriers and normal for non-carriers. Moreover, through manual inspection, we identified a region within the *KITLG* gene near an exon with a normalized coverage difference of 1.14 (Chr5:94,081,500-94,081,750; Additional File 13 Figure S6). A detailed view of this region in Integrative Genomics Viewer (Additional File 14 Figure S7) indicates a potential insertion at this location, linked to the haplotype we identified.

A more exhaustive analysis of structural variants was conducted in eighteen pigs for which HiFi long-read sequencing data were available. From these eighteen pigs, six were heterozygous carriers and one was homozygous for the alternate allele of the missense variant 5_94084790_G>A. An alignment-based approach identified 445 structural variants in the interval from 92 Mb to 96 Mb on SSC5 including sixteen that were in high LD (r2>0.8) with the missense variant 5_94084790_G>A (Additional File 15 Table S8 and Additional File 16 Figure S8). There were no SVs overlapping coding sequence, but we identified a 316 bp insertion at Chr5:94,081,687 bp in the third intron of *KITLG* (317 bp upstream of exon 4; Additional File 15 Table S8 and Additional File 16 Figure S8). This insertion matches the repeat annotation for the porcine repetitive element 1 (PRE1), a short interspersed nuclear element (SINE) [38]. This insertion at Chr5:94,081,687 bp and the deletion at Chr5: 94,051,620 bp also coincides with the region detected by manually scanning the *KITLG* region in IGV for the 120 short-read sequenced individuals (Additional File 14 Figure S7). There was also a 288 bp deletion at Chr5:94,014,953, 2.4 Kb upstream of the first *KITLG* exon (Additional File 15 Table S8, Additional File 13 Figure S8). This deletion removes a putative Pre0_SS element, which is also a SINE element of the PRE1 family [39] and is just outside a region with a normalized coverage difference greater than three in the short read data (Chr5:94,015,000-94,015,250; Additional File 13 Figure S6). The 316 bp insertion at Chr5:94,081,687 bp and the 288 bp deletion at Chr5:94,014,953 were in perfect LD with the missense variant 5_94084790_G>A.

## Discussion

In this study, we investigated the prevalence and genetics of iris pigmentation in the Swiss Landrace and Swiss Large White pig populations. We found a remarkably high prevalence for heterochromia iridum (18.6%) in the Swiss Landrace breed. Iris pigmentation was highly heritable (h^2^=56.0-57.2%) and it was not correlated with production traits. Our GWAS identified multiple significant associations near candidate genes such as *TYR*, *ALX4, DCT* and *KITLG*. Fine-mapping identified a missense variant (5_94084790_G>A) as a plausible candidate causative variant for pale and heterochromatic iris pigmentation in Swiss pigs. However, further functional investigations are needed to establish causality of the missense variant or other candidate causal variants.

The observed prevalences of iris pigmentation phenotypes in the Swiss breeds were comparable to those reported previously in other populations (see references in Table 1). However, direct comparisons need to be interpreted with caution due to the limited number of available studies and substantial differences in phenotyping methods, populations examined, and time periods. Iris pigmentation variability in the Swiss Large White population was comparable to that reported in Italian Large White pigs [4], phenotyped with a similar methodology. Nonetheless, we observed a somewhat higher prevalence of pale irises (6.4% vs 3.8%), heterochromia iridum (7.0% vs 5.9%), and heterochromia iridis (10.0% vs 3.2%). The most striking observation was the 18.6% prevalence of heterochromia iridum in Swiss Landrace pigs, which is higher than previously reported in other, non-Landrace populations (see references in Table 1). Heterochromia iridum is very rare (<1%) in humans [40]. Our findings of a high inter-breed variability of heterochromia iridum in pigs is in line with observations in dogs and cats, where this trait has remarkably high prevalences in some breeds (e.g., Siberian Husky, Border Collie) but may be much rarer in others [41]. As cats and dogs are companion animals, they may have been actively selected for iris pigmentation, such as pale or heterochromatic irises [42]. However, active selection for specific iris pigmentation phenotypes seems unlikely in Swiss Landrace pigs, as iris pigmentation is often obscured by their overhanging ears and no significant genetic correlations with major production traits were identified. Instead, the high incidence of heterochromia may be explained by genetic drift and/or genetic hitchhiking linked to other (re)productive traits not examined in this study.

Iris pigmentation is highly heritable (h² = 56.0–57.2%) on the liability scale in the two pig populations studied. While this is consistent with heritability estimates in humans (h² = 51%, H² = 85%) [8], our estimates are higher than those reported previously in other pig populations (h² = 9.5–50.4%) [4]. This discrepancy may be partly due to differences in the statistical approaches used to estimate heritability. We employed a categorical threshold model that integrated both pedigree and genomic information. This method might be better to account for additive genetic effects and familial structure, whereas Moscatelli et al. [4] relied solely on SNP-based data analyzed with the GEMMA software under various iris pigmentation scenarios.

Our GWAS yielded several associations that were significant at the genome-wide significance threshold (P<5×10^-8^). The strongest associations across breeds were found on SSC5 near the gene *KIT ligand* (*KITLG;* Chr5:94,017,387–94,110,214). The *KITLG* gene is a pleiotropic cytokine that binds the KIT receptor to regulate diverse processes including melanogenesis, hematopoiesis, stem cell maintenance, and cell migration, acting through pathways like PI3K-AKT, MAPK, and STAT signaling [11]. *KITLG* has been associated with pigmentation traits in several species, such as pigs [4,5], cattle [43], goats [44], and humans [13–15,45].

Our fine-mapping efforts prioritized a missense variant (5_94084790_G>A) in the *KITLG* gene as a candidate causal variant for pale and heterochromatic irises in pigs. The prioritized missense variant is close to the top variant reported by Moscatelli et al. [4] (Chr5:94,284,630; P=1.0×10^-11^).

Several *KITLG* missense variants affecting the *KIT* binding domain have been identified previously to underly pigmentation disorders in humans [12,45] including a loss-of-function variant (p.Leu104Val) which also causes heterochromia iridis and progressive hearing loss [15]. The missense variant associated with porcine eye pigmentation also resides in the KIT binding domain of *KITLG*. We did not recognize skin hypo- or hyperpigmentation in the pigs we phenotyped and their hearing ability was not systematically evaluated. Behavioral abnormalities that could possibly result from impaired hearing were not reported by the farmers. It is therefore likely that the prioritized missense variant is not a loss-of-function allele which agrees with a SIFT score of 1 (range of 0–1, where 1 is benign and 0 is deleterious) which suggests it is benign for protein function.

Several small and structural variants are in LD with the missense variant but none of them was a compelling candidate although some overlapped putative regulatory elements. A possible impact of structural variations on expression and function of *KITLG* has been described in previous research. In dogs, for example, a 6 kb copy number variant upstream of *KITLG* has been associated with pigment intensity of the coat [46], and Guenther et al. [47] showed in mice that a regulatory enhancer of *KITLG* influenced hair pigmentation. Further research involving functional data is required to establish causal links between the missense variant and any other highly significantly associated variants identified nearby and iris pigmentation.

Our GWAS also identified several associations surpassing the suggestive threshold (P<10^-5^, Additional File 10 Table S5) nearby genes that have been implicated to underly eye, skin, or coat color variation in other species. The close vicinity of functional candidate genes adds confidence that some of the suggestive associations are true positives. For instance, we found suggestive associations nearby *KIT*, *DCT*, *ALX4*, *SFMBT2* and *TYR [5,7,9,41,48–50]*. The associations in the Swiss Landrace and Swiss Large White breeds near *ALX4 (*Chr2:18,037,401-18,085,683) are noteworthy considering that variants nearby *ALX4* were strongly associated with pale irises and heterochromia iridum in Siberian Huskies [41].

This study presents novel insights into the genetic basis of iris pigmentation in pigs. However, several limitations should be acknowledged. First, phenotyping was conducted under practical field conditions on live animals, which may have introduced inaccuracies, particularly in Swiss Landrace pigs where overhanging ears often obscure the eyes. This may have led to under- or misclassification of iris pigmentation phenotypes, especially subtle variations such as light versus medium brown irises. Future studies could benefit from (post-mortem) assessment using standardized tools such as reflectance spectrometry or high-resolution imaging to improve accuracy and reproducibility of pigmentation scoring. Second, although whole-genome sequence data enabled the detection of promising candidate variants and structural changes near the *KITLG* gene, these findings were not directly supported by animals with phenotypic records and sequence data. This limitation restricts the ability to definitively link specific mutations to observed phenotypes. Expanding the dataset to include phenotyped individuals with matched high-resolution genotypes or sequence information would enhance the power to confirm causal variants. Third, the quality of SNP imputation may have been suboptimal, particularly for the Swiss Landrace population, which was imputed using the publicly available SWIM tool [24]. Although the SWIM tool contains 651 Landrace animals in the reference panel, these pigs come from different populations than the Swiss Landrace pigs. These differences in population structure or reference panel composition may have impacted imputation accuracy, potentially affected downstream association results, and limited power. Fourth, this study did not include functional validation of the identified candidate variants near *KITLG*. Without experimental follow-up, these variants remain putative. While phenotyping and data limitations warrant cautious interpretation, this work provides a strong foundation for future functional studies on pigmentation traits in pigs.

## Conclusion

The present study provides new insights into the genetic architecture of iris pigmentation in pigs. We report a remarkably high prevalence of heterochromia iridum in Swiss Landrace pigs (18.6%) and demonstrate that iris pigmentation is a highly heritable trait (h² = 56.0–57.2%) that is genetically not significantly correlated to production traits. Through genome-wide association analyses, we identified several candidate loci associated with iris pigmentation, with the strongest and most consistent signals near the *KITLG* gene on SSC5. Our data support the involvement of a missense variant (5_94084790_G>A) as potential causal mutation underlying pale and heterochromatic irises. Moreover, several structural variants, such as a PRE1 insertion, are in high linkage disequilibrium with this missense variant in Swiss Large White pigs. Additional associations near genes such as *DCT*, *ALX4*, *TYR* and *KIT* suggest a polygenic inheritance of porcine eye colour involving multiple pigmentation-related pathways. While phenotyping and data limitations warrant cautious interpretation, this work lays a strong foundation for future functional studies on pigmentation traits in pigs.

## Supporting information

Summary_Additional_Files

## List of abbreviations

GWAS: Genome-wide association study
H^2^: Broad-sense heritability
h^2^: Narrow-sense heritability
HPD: 95% highest posterior density distribution
IGV: Integrative Genomics Viewer
R_g_: Genetic correlation
N: Number of individuals, sample size
PCA: Principal component analysis
SINE: Short interspersed nuclear element
SSC: *sus scrofa* chromosome
VEP: Variant effect predictor

## Declarations

## Ethics approval and consent to participate

The pigs described in this study were kept in compliance with the Swiss legislations and were not raised or treated in any way for the purpose of this study. Genotype and sequence data were provided by SUISAG and collected within their breeding program. Therefore, ethical approval was not necessary for this study.

## Consent for publication

Not applicable

## Availability of data and materials

Genotyping data were provided by SUISAG. The datasets generated and/or analyzed during the current study are not publicly available due to restrictions from SUISAG.

High-coverage sequencing read data of the 120 Large White pigs used in this study are available at the European Nucleotide Archive (ENA) (http://www.ebi.ac.uk/ena) of the EMBL at BioProject PRJEB38156 and PRJEB39374.

## Authors’ contributions

WG analysed the data and wrote the manuscript. NK helped in collecting the data and was responsible for data transfer. NKK developed computational workflows that were used in this study, e.g., for haplotype-based association analysis. AM and ASL performed long read and structural variant analysis. WG and HP designed and conceived this study. SN contributed to genomic data collection. WG, NKK, NK, ASL, QH, AM, SN and HP critically reviewed the analyses and the manuscript. All authors have read and approved the final manuscript.

## Acknowledgements

We would like to thank all SUISAG employees and farmers that collected the data for this study. We acknowledge SUISAG for all their help and sharing their data. Thanks to Nicole Maffezzini for her contribution.

## Additional files

### Additional File 1 Figure S1

Format: .jpg

Title: Phenotyping scale for iris pigmentation, following Moscatelli et al. (2020)

Description: Phenotyping scale for iris pigmentation, following Moscatelli et al. (2020). In this study, medium and dark brown irises were grouped together in ‘dark’, as it was hard to distinguish both types in live pigs in a farm environment.

### Additional File 2 Figure S2

Format: .jpg

Title: Histogram of production traits distribution for the Swiss Landrace population.

Description: Top left: Histogram of the age at recording phenotype. Top right: Histogram of life daily gain (g/d). Bottom left: Histogram of backfat thickness (mm). Bottom right: Histogram of loin muscle depth (mm).

### Additional File 3 Figure S3

Format: .jpg

Title: Histogram of production traits distribution for the Swiss Large White population.

Description: Top left: Histogram of the age at recording phenotype. Top right: Histogram of life daily gain (g/d). Bottom left: Histogram of backfat thickness (mm). Bottom right: Histogram of loin muscle depth (mm).

### Additional File 4 Figure S4

Format: .png

Title: Principal component (PC) analysis of the Swiss Landrace and Swiss Large White array-derived genotypes.

Description: Left: PC1 versus PC2, showing different subpopulations. Right: PC1 versus PC3, showing Swiss Large White subpopulations cluster together.

### Additional File 5 Figure S5

Format: .png

Title: Principal component (PC) analysis of genotyped Swiss Large White pigs versus sequenced Swiss Large White pigs.

Description: Left: PC1 versus PC2 Right: PC1 versus PC3

### Additional File 6 Table S1

Format: .xlsx

Title: Overview of number of phenotyped pigs per farm and per breed

Description: Number of pigs with a specified iris pigmentation phenotype per farm and per breed. In total, there were 837 Swiss Landrace, 328 Swiss Large White, 20 crossbreds from Swiss Landrace and Swiss Large White, 11 Duroc and 4 Piétrain pigs phenotyped. LDR: Swiss Landrace; LWT: Swiss Large White; HYB: Hybrid sow crossbred from Swiss Landrace and Swiss Large White; DUR: Duroc; PIT: Piétrain

### Additional File 7 Table S2

Format: .xlsx

Title: Prevalence of offspring iris pigmentation phenotypes versus maternal phenotype for Swiss Landrace.

Description: The number of phenotyped sows (N_sow_) and the number of phenotyped offspring (N_off_) per sow phenotype are given between brackets. Heterochromia iridis unilateralis and bilateralis were combined to have a relevant sample size.

### Additional File 8 Table S3

Format: .xlsx

Title: Genetic parameters and heritability estimates for iris pigmentation in single trait 5-category threshold model.

Description: Main genetic parameters are shown for the single trait analysis for both the Swiss Landrace and the Swiss Large White.

### Additional File 9 Table S4

Format: .xlsx

Title: Genetic parameters, heritability estimates and genetic correlations estimated for iris pigmentation and production traits in a bivariate threshold model with a 5-category trait and a linear trait.

Description: Main genetic parameters are shown for the bivariate analysis for both the Swiss Landrace and the Swiss Large White. Swiss Large White estimates were not shown in the manuscript due to a limited sample size.

### Additional File 10 Table S5

Format: .xlsx

Title: GWAS results of most significant associations per 1Mb bin per scenario surpassing the suggestive threshold of P<10^-5^.

Description: GWAS results of most significant associations per 1Mb bin per scenario surpassing the suggestive threshold of P<10^-5^ for the three studied populations (‘breed’; Swiss Landrace, Swiss Large White and combined) and GWAS methods (haplotypes or imputed sequences) for different scenarios as explained in Table 3 (‘trait’).

### Additional File 11 Table S6

Format: .xlsx

Title: GWAS results of all variants surpassing the suggestive threshold of P<10^-5^.

Description: GWAS results of all variants surpassing the suggestive threshold of P<10^-5^ for the three studied populations (‘breed’; Swiss Landrace, Swiss Large White and combined) and GWAS methods (haplotypes or imputed sequences) for different scenarios as explained in Table 3 (‘trait’).

### Additional File 12 Table S7

Format: .xlsx

Title: VEP annotation of 14,835 iris pigmentation-associated variants

Description: This table contains the Ensembl Variant Effect Predictor (VEP) annotations for 14,835 SNPs and indels that surpassed the suggestive genome-wide association threshold (P < 1×10⁻⁵) in at least one iris pigmentation GWAS comparison. Each variant is annotated with its predicted functional consequence, including coding impact (e.g., synonymous, missense), location (e.g., intronic, intergenic), and overlap with known genes or regulatory features.

### Additional File 13 Figure S6

Format: .png

Title: Mean normalized coverage per haplotype associated with iris pigmentation in Swiss Large White pigs

Description: Mean normalized coverage was estimated via Mosdepth per 250 bp interval based on short-read sequence data of 120 Large White pigs with at least 10x coverage. From this cohort, 6 homozygous carriers were identified for a haplotype associated with iris pigmentation, 39 heterozygous carriers and 75 non-carriers. Left: Mean normalized coverage distance within the *KITLG* gene near exon 2. Right: Mean normalized coverage distance near a 288bp deletion at Chr5:94,014,953, just 2.4 Kb upstream of the first *KITLG* exon (Additional File 15 Table S8, Additional File 16 Figure S8). This deletion matched the Pre0_SS element, which is also a SINE element of the PRE1 family.

### Additional File 14 Figure S7

Format: .png

Title: Integrative Genomics Viewer (IGV) plot showing a potential insertion in the six homozygous haplotype carriers (top tracks) compared to randomly selected non-carriers (bottom tracks).

Description: The plot displays read alignments and genome coverage at chromosome 5 (Chr5:94,081,200–94,082,100), with a suspected insertion site around Chr5:94,081,500. This region is located near the second exon of the KITLG gene.

### Additional File 15 Table S8

Format: .xlsx

Title: Insertions and deletions detected near the *KITLG* gene in LD (r^2^>0.8) with missense variant 5_94084790_G>A.

Description: Structural variants in high linkage disequilibrium (LD; r2>0.8) with the missense variant 5_94084790_G>A in the 92-96Mb region near the KITLG on SSC5.

### Additional File 16 Figure S8

Format: .png

Title: Insertions and deletions detected near the *KITLG* gene in LD (r^2^>0.8) with missense variant 5_94084790_G>A.

Description: For the 18 pigs with HiFi long-read sequence information, several deletions and insertions were found in LD (r^2^>0.8) with missense variant 5_94084790_G>A.

