## Supplementary material for "Genetic analysis of iris pigmentation in Swiss pig breeds identifies a missense *KITLG* variant as a potential causal factor for pale and heterochromatic irises": Summary_Additional_Files

*8092 Zürich, Switzerland*

[2]*SUISAG, Allmend 10, Sempach, 6204, Switzerland*

[3]Animal Genetics unit, Tannenstrasse 1, 8092 Zürich

[$] *current affiliation: Generatio GmbH, Heidelberg, Germany 69126*

*Corresponding author

E-mail addresses:

WG:

NKK:

NK:

ASL:

QH:

AM:

SN:

HP:

### Additional files

**Additional File 1 Figure S1**

Format: .jpg

Title: Phenotyping scale for iris pigmentation, following Moscatelli et al. (2020)

Description: Phenotyping scale for iris pigmentation, following Moscatelli et al. (2020). In this study, medium and dark brown irises were grouped together in ‘dark’, as it was hard to distinguish both types in live pigs in a farm environment.

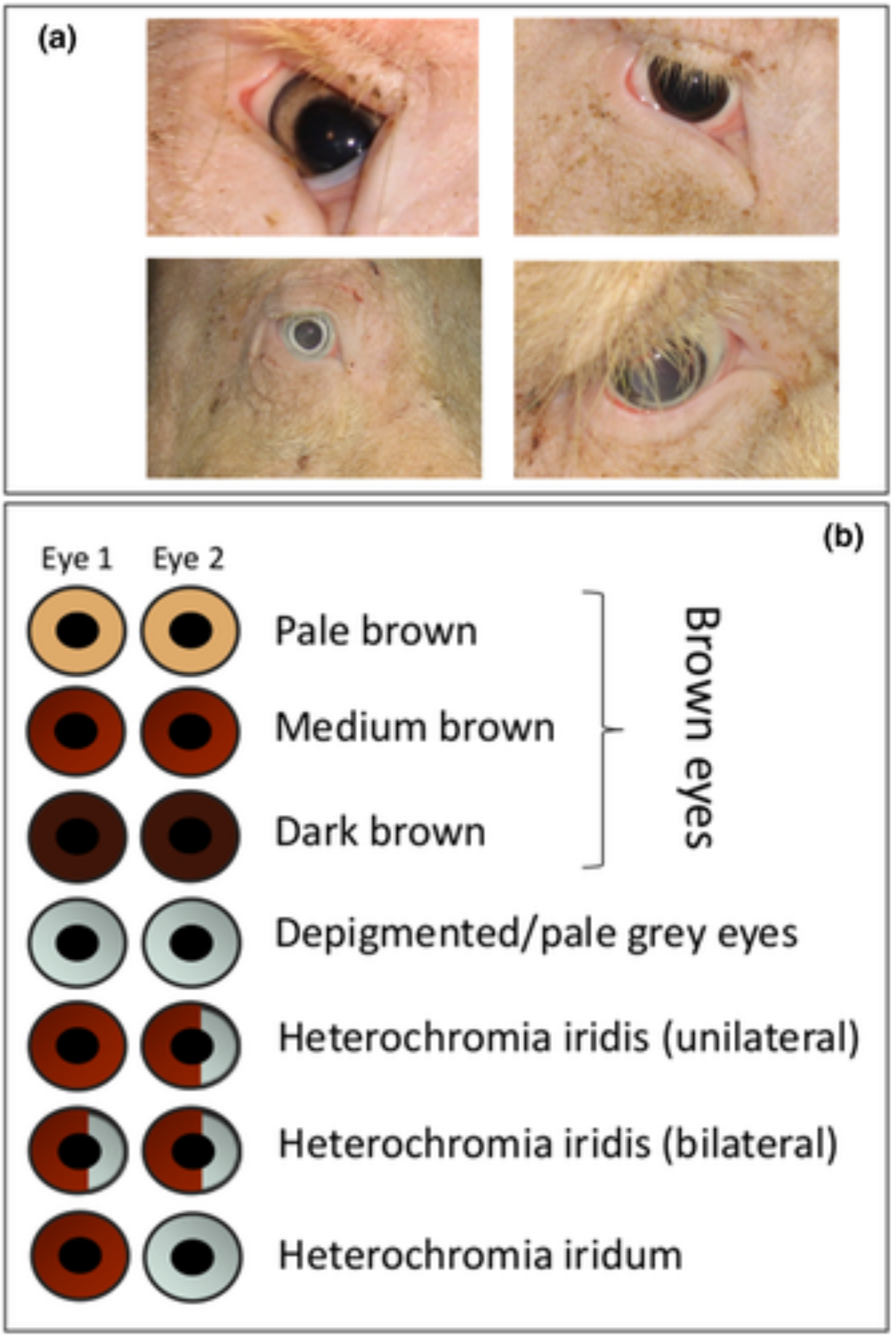

**Additional File 2 Figure S2**

Format: .jpg

Title: Histogram of production traits distribution for the Swiss Landrace population.

Description: Top left: Histogram of the age at recording phenotype. Top right: Histogram of life daily gain (g/d). Bottom left: Histogram of backfat thickness (mm). Bottom right: Histogram of loin muscle depth (mm).

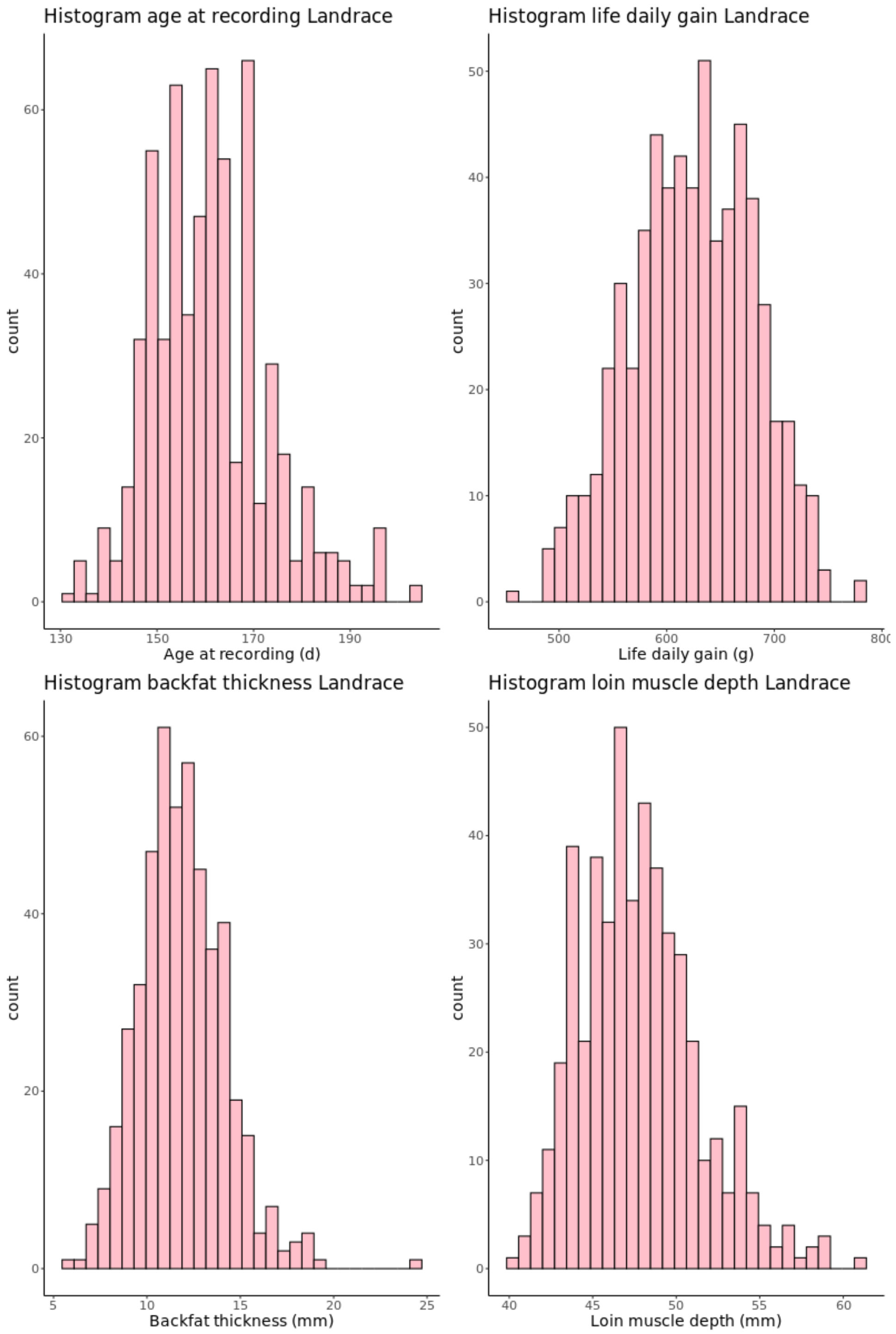

**Additional File 3 Figure S3**

Format: .jpg

Title: Histogram of production traits distribution for the Swiss Large White population.

Description: Top left: Histogram of the age at recording phenotype. Top right: Histogram of life daily gain (g/d). Bottom left: Histogram of backfat thickness (mm). Bottom right: Histogram of loin muscle depth (mm).

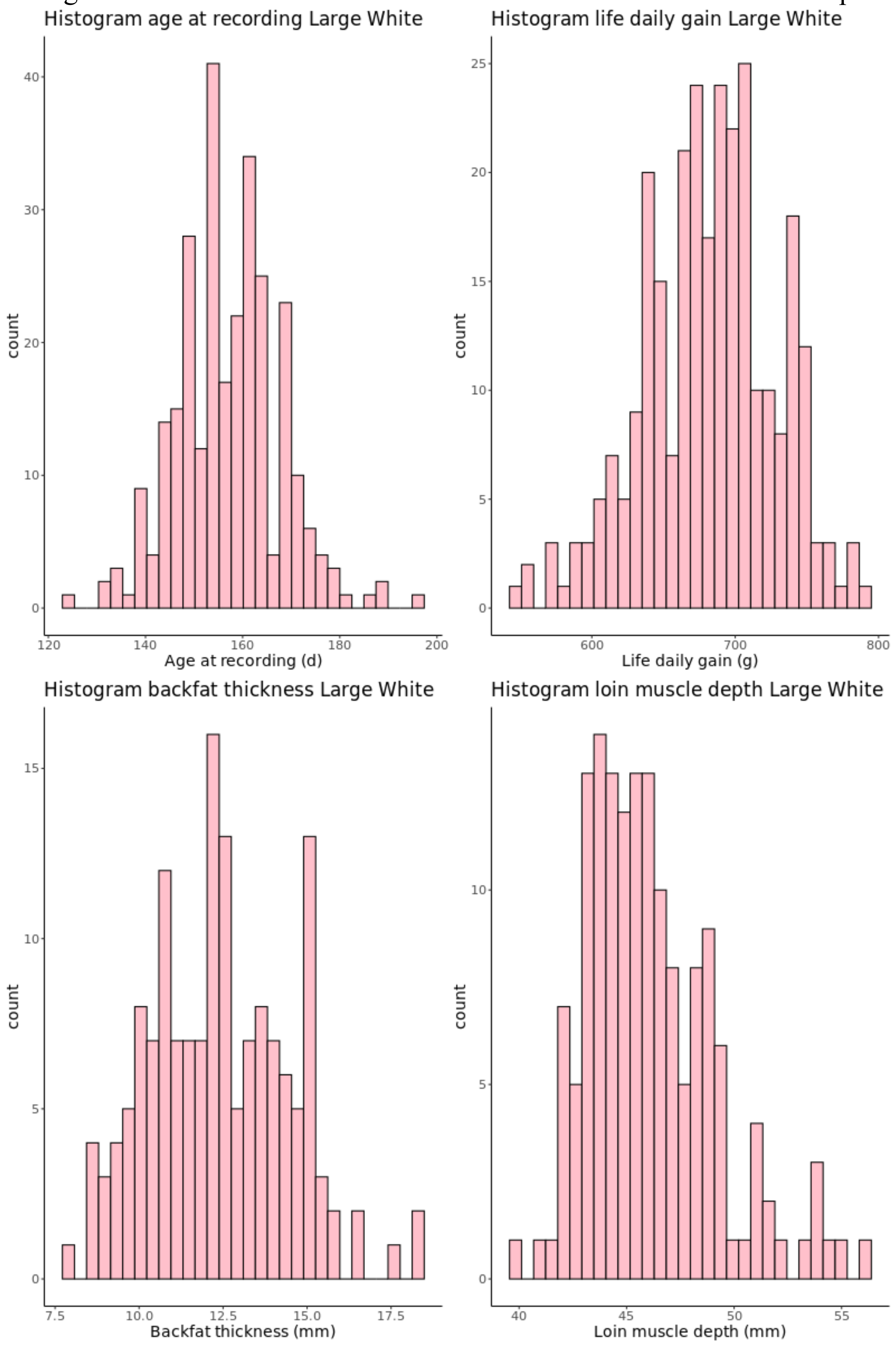

**Additional File 4 Figure S4**

Format: .png

Title: Principal component (PC) analysis of the Swiss Landrace and Swiss Large White array-derived genotypes.

Description: Left: PC1 versus PC2, showing different subpopulations. Right: PC1 versus PC3, showing Swiss Large White subpopulations cluster together.

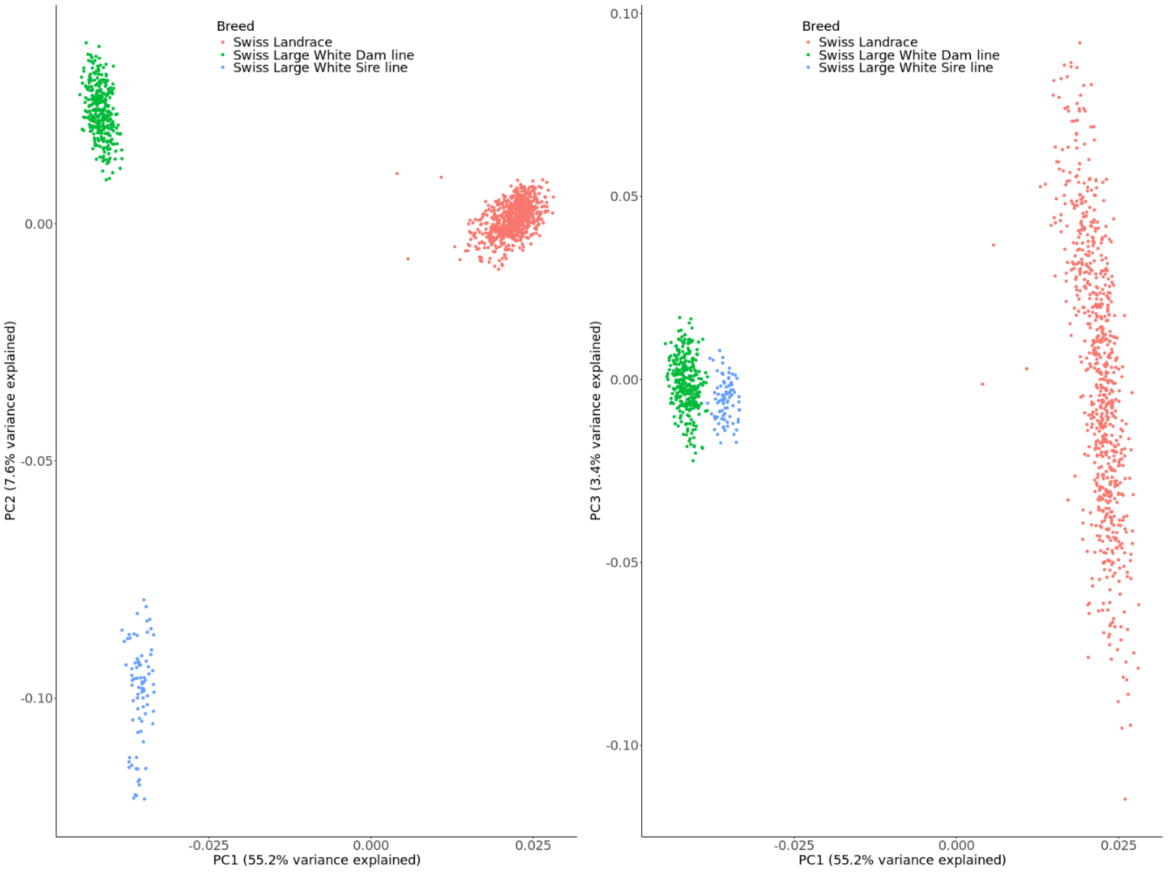

**Additional File 5 Figure S5**

Format: .png

Title: Principal component (PC) analysis of genotyped Swiss Large White pigs versus sequenced Swiss Large White pigs.

Description: Left: PC1 versus PC2 Right: PC1 versus PC3

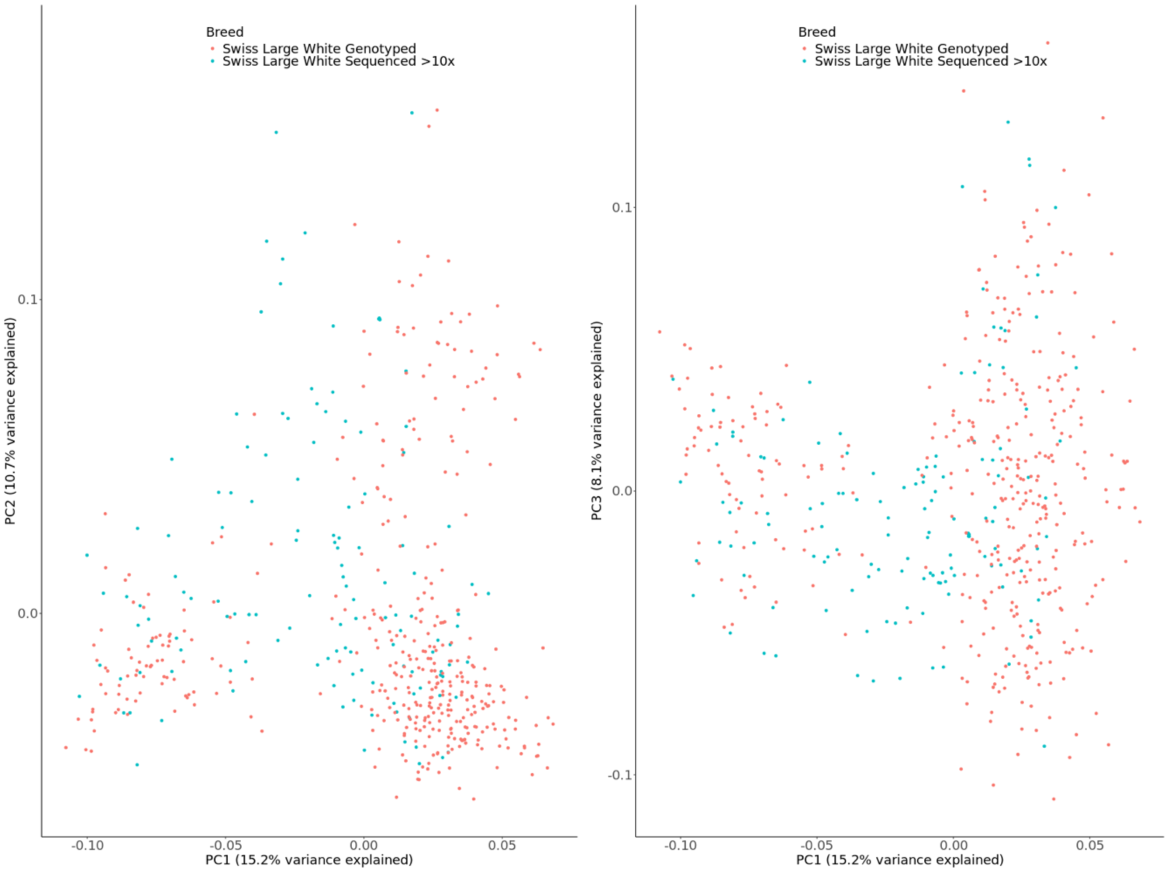

**Additional File 6 Table S1**

Format: .xlsx

Title: Overview of number of phenotyped pigs per farm and per breed

Description: Number of pigs with a specified iris pigmentation phenotype per farm and per breed. In total, there were 837 Swiss Landrace, 328 Swiss Large White, 20 crossbreds from Swiss Landrace and Swiss Large White, 11 Duroc and 4 Piétrain pigs phenotyped. LDR: Swiss Landrace; LWT: Swiss Large White; HYB: Hybrid sow crossbred from Swiss Landrace and Swiss Large White; DUR: Duroc; PIT: Piétrain

| | **Farm A** | | **Farm B** | | **Farm C** | **Farm D** | | **Farm E** | | **SUISAG testing station** | | | |
|---|---|---|---|---|---|---|---|---|---|---|---|---|---|
| **Breed** | **LDR** | **LWT** | **LDR** | **HYB** | **LDR** | **LDR** | **LWT** | **LDR** | **LWT** | **LDR** | **LWT** | **PIT** | **DUR** |
| **Total** | **127** | **11** | **125** | **20** | **58** | **98** | **50** | **158** | **4** | **271** | **263** | **4** | **11** |
| **Dark Brown** | 61 | 7 | 77 | 15 | 29 | 39 | 22 | 76 | 2 | 125 | 135 | 1 | 3 |
| **Light Brown** | 11 | 2 | 19 | 3 | 9 | 17 | 8 | 28 | 2 | 53 | 73 | 4 | 8 |
| **Pale** | 15 | 0 | 8 | 0 | 4 | 9 | 9 | 8 | 0 | 17 | 12 | 0 | 0 |
| **Heterochromia iridis unilateralis** | 13 | 1 | 4 | 1 | 3 | 10 | 7 | 7 | 0 | 27 | 20 | 0 | 0 |
| **Heterochromia iridis bilateralis** | 2 | 0 | 0 | 0 | 0 | 2 | 1 | 2 | 0 | 6 | 4 | 0 | 0 |
| **Heterochromia iridum** | 25 | 1 | 17 | 1 | 13 | 21 | 3 | 37 | 0 | 43 | 19 | 0 | 0 |

**Additional File 7 Table S2**

Format: .xlsx

Title: Prevalence of offspring iris pigmentation phenotypes versus maternal phenotype for Swiss Landrace.

Description: The number of phenotyped sows ($N_{sow}$) and the number of phenotyped offspring ($N_{off}$) per sow phenotype are given between brackets. Heterochromia iridis unilateralis and bilateralis were combined to have a relevant sample size.

| | Iris pigmentation sow | | | | |
|---|---|---|---|---|---|
| **Iris pigmentation offspring** | **Dark brown** ($N_{sow}$=69; $N_{off}$=142) | **Light brown** ($N_{sow}$=20; $N_{off}$=69) | **Pale** ($N_{sow}$=13; $N_{off}$=34) | **Iridum** ($N_{sow}$=24; $N_{off}$=72) | **Iridis** ($N_{sow}$=12; $N_{off}$=42) |
| **Dark Brown** | 64.1% | 55.1% | 38.2% | 36.1% | 52.4% |
| **Light Brown** | 10.6% | 10.1% | 14.7% | 19.4% | 19.1% |
| **Pale** | 4.9% | 10.1% | 5.9% | 4.2% | 4.8% |
| **Iridum** | 14.8% | 20.3% | 23.5% | 25.0% | 19.1% |
| **Iridis** | 5.6% | 4.4% | 17.7% | 15.3% | 4.8% |

**Additional File 8 Table S3**

Format: .xlsx

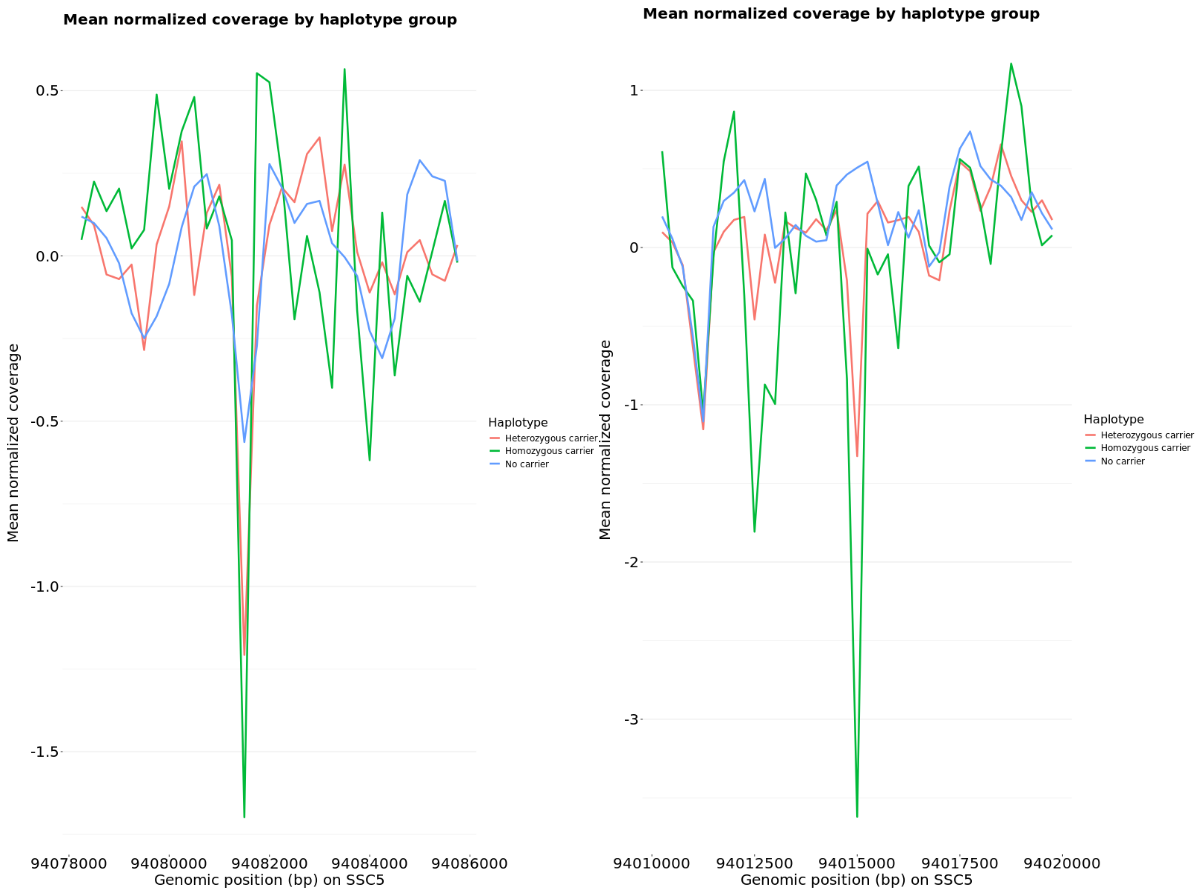

**Additional File 14 Figure S7**

Format: .png

Title: Integrative Genomics Viewer (IGV) plot showing a potential insertion in the six homozygous haplotype carriers (top tracks) compared to randomly selected non-carriers (bottom tracks).

Description: The plot displays read alignments and genome coverage at chromosome 5 (Chr5:94,081,200–94,082,100), with a suspected insertion site around Chr5:94,081,500. This region is located near the second exon of the KITLG gene.

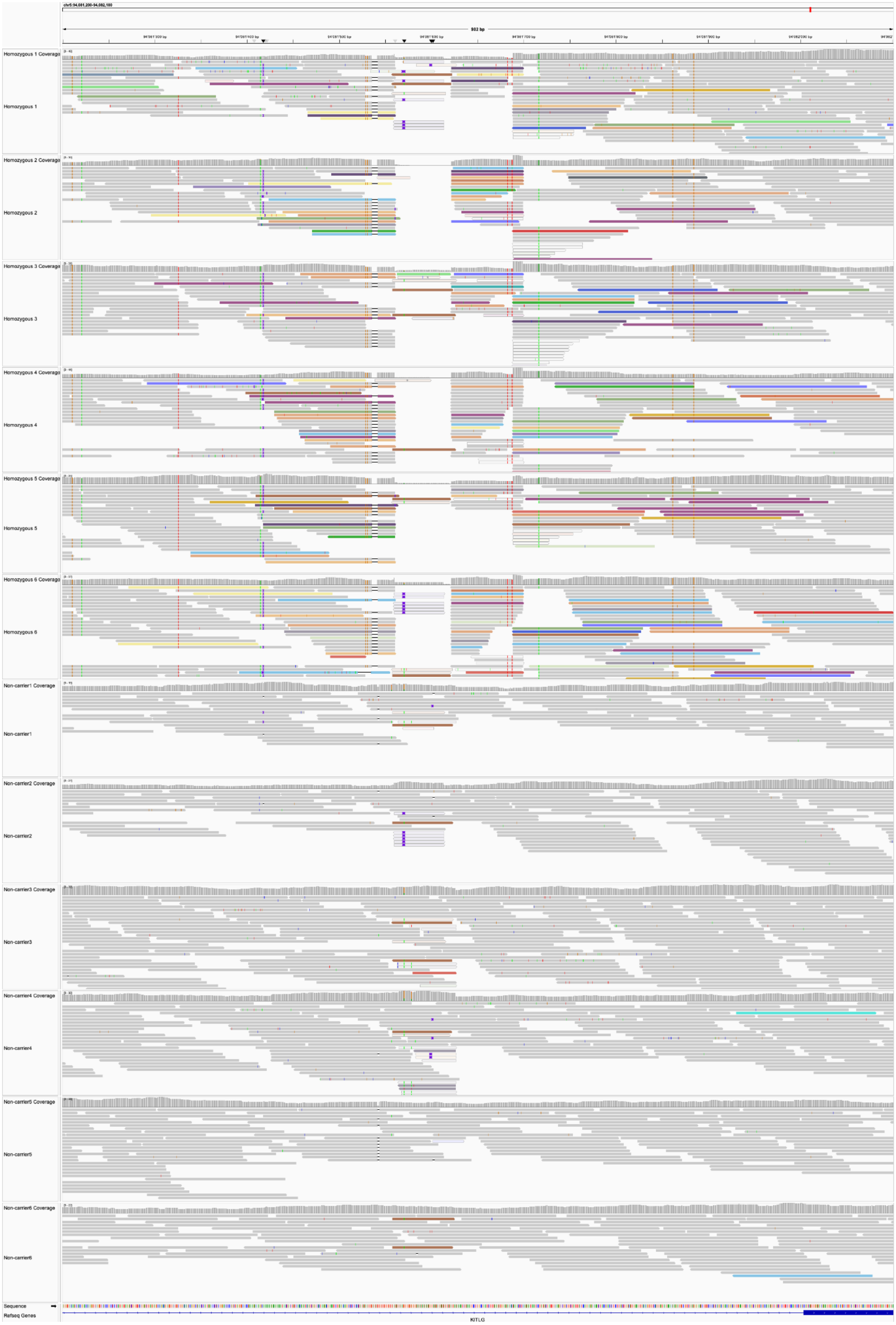
chr5:94,081,200-94,082,100
902 bp
94'081'300 bp
94'081'400 bp
94'081'500 bp
94'081'600 bp
94'081'700 bp
94'081'800 bp
94'081'900 bp
94'082'000 bp
Homozygous 1 Coverage
Homozygous 1
Homozygous 2 Coverage
Homozygous 2
Homozygous 3 Coverage
Homozygous 3
Homozygous 4 Coverage
Homozygous 4
Homozygous 5 Coverage
Homozygous 5
Homozygous 6 Coverage
Homozygous 6
Non-carrier1 Coverage
Non-carrier1
Non-carrier2 Coverage
Non-carrier2
Non-carrier3 Coverage
Non-carrier3
Non-carrier4 Coverage
Non-carrier4
Non-carrier5 Coverage
Non-carrier5
Non-carrier6 Coverage
Non-carrier6
Sequence
Refseq Genes
KITLG

**Additional File 15 Table S8**

Format: .xlsx

Title: Insertions and deletions detected near the *KITLG* gene in LD ($r^2$>0.8) with missense variant 5_94084790_G>A.

Description: Structural variants in high linkage disequilibrium (LD; r2>0.8) with the missense variant 5_94084790_G>A in the 92-96Mb region near the KITLG on SSC5.

**Additional File 16 Figure S8**

Format: .png

Title: Insertions and deletions detected near the *KITLG* gene in LD ($r^2>0.8$) with missense variant 5_94084790_G>A.

Description: For the 18 pigs with HiFi long-read sequence information, several deletions and insertions were found in LD ($r^2>0.8$) with missense variant 5_94084790_G>A.

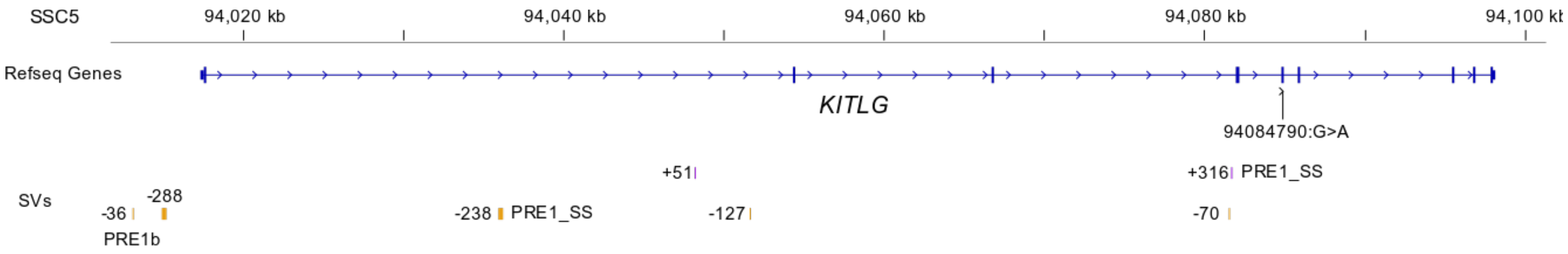
